# Oropouche virus glycoprotein topology and cellular requirements for virus assembly

**DOI:** 10.1101/2020.05.28.122689

**Authors:** Natalia S. Barbosa, Juan O. Concha, Luis L. P. daSilva, Colin M. Crump, Stephen C. Graham

**Affiliations:** Center for Virus Research, Ribeirão Preto Medical School, University of São Paulo, Ribeirão Preto, SP, Brazil; Department of Cell and Molecular Biology, Ribeirão Preto Medical School, University of São Paulo, Ribeirão Preto, SP, Brazil; Department of Pathology, University of Cambridge, Cambridge, United Kingdom

**Keywords:** Bunyavirus, Bunyamwera virus, polyprotein processing, virus budding

## Abstract

Oropouche virus (OROV; *Genus: Orthobunyavirus*) is the etiological agent of Oropouche fever, a debilitating febrile illness common in South America. We used recombinant expression of the OROV M polyprotein, that encodes the surface glycoproteins Gn and Gc plus the non-structural protein NSm, to probe the cellular determinants for OROV assembly and budding. Gn and Gc self-assemble and are secreted independently of NSm. Mature OROV Gn has two predicted transmembrane domains that are crucial for glycoprotein translocation to the Golgi complex and glycoprotein secretion and, unlike related orthobunyaviruses, both transmembrane domains are retained during Gn maturation. Disruption of Golgi function using the drugs brefeldin A and monensin inhibit glycoprotein secretion. Infection studies have previously shown that the cellular Endosomal Sorting Complexes Required for Transport (ESCRT) machinery is recruited to Golgi membranes during OROV assembly and that ESCRT activity is required for virus secretion. A dominant negative form of the ESCRT-associated ATPase VPS4 significantly reduces recombinant OROV glycoprotein secretion and blocks virus release from infected cells, and VPS4 partly co-localizes with OROV glycoproteins and membranes co-stained with Golgi markers. Furthermore, immunoprecipitation and fluorescence microscopy experiments demonstrate that OROV glycoproteins interact with the ESCRT-III component CHMP6, with overexpression of a dominant negative form of CHMP6 significantly reducing OROV glycoprotein secretion. Taken together, our data highlights differences in M polyprotein processing across orthobunyaviruses, that Golgi and ESCRT function are required for glycoprotein secretion, and identifies CHMP6 as an ESCRT-III component that interacts with OROV glycoproteins.

**Importance:** Oropouche virus causes Oropouche fever, a debilitating illness common in South America that is characterised by high fever, headache, myalgia and vomiting. The tripartite genome of this zoonotic virus is capable of reassortment and there have been multiple epidemics of Oropouche fever in South America over the last 50 years, making Oropouche virus infection a significant threat to public health. However, the molecular characteristics of this arbovirus are poorly understood. We developed a recombinant protein expression system to investigate the cellular determinants of OROV glycoprotein maturation and secretion. We show that the proteolytic processing of the M polypeptide, which encodes the surface glycoproteins (Gn and Gc) plus a non-structural protein (NSm), differs between OROV and its close relative Bunyamwera virus. Furthermore, we demonstrate that OROV M glycoprotein secretion requires the cellular ESCRT membrane-remodelling machinery and identify that the OROV glycoproteins interact with the ESCRT protein CHMP6.

## Introduction

Oropouche virus (OROV) is an arbovirus that is the etiological agent of Oropouche fever, a debilitating febrile illness. Oropouche fever symptoms range from high fever to vomiting, photophobia and, in rare cases, aseptic meningitis or meningoencephalitis (1). OROV is prevalent in the Caribbean and tropical regions of Latin America, and more than 30 epidemics of Oropouche fever have occurred since the first isolation of OROV in 1955 (1). Clinical diagnosis of Oropouche fever is challenging due to the resemblance of its symptoms to diseases caused by other arboviruses like Dengue virus, Zika virus and Chikungunya virus (2). There are no specific antiviral treatments for Oropouche fever, nor is there an effective vaccine to prevent OROV infection. The zoonotic origin, history of human spill-over, tri-segmented RNA genome that is capable of re-assortment (3) and increasing contact between humans and wild-animal reservoirs due to deforestation (1) make OROV a serious epidemic threat.

OROV belongs to *Orthobunyavirus* genus of the *Peribunyaviridae* family, one of the fourteen families of the order *Bunyavirales*. Orthobunyaviruses form spherical enveloped virus particles that are 100– 120 nm in diameter and orthobunyavirus genomes comprise three negative sense single-stranded RNA segments: The small segment (S) encodes the nucleocapsid N protein (25–30 kDa), which oligomerizes and encapsidates the viral genome, and the non-structural protein NSs; The medium segment (M) encodes a polyprotein that is co-translationally cleaved by host proteases to yield the viral surface glycoproteins (Gc and Gn) plus a non-structural protein NSm; The large segment (L) encodes the viral RNA dependent RNA polymerase (RdRp) that catalyzes viral replication and transcription (4). The viral RNA segments are encapsidated by N protein to form a ribonucleoprotein complex that associates with both the RdRp and the surface glycoproteins to promote virus particle assembly (5). The Gn (∼32 kDa) and Gc (∼110 kDa) glycoproteins are integral membrane proteins with N-terminal ectodomains and they associate in the host endoplasmic reticulum (ER) before being transported to the Golgi complex, the main site of virion assembly (6–8). Studies with Bunyamwera virus (BUNV), the prototypical orthobunyavirus, have shown that the virus particles undergo distinct morphological changes inside Golgi and *trans*-Golgi network cisternae (7), and studies of both BUNV and OROV have shown profound fragmentation on Golgi complex cisternae during virus infection of some cell types (6, 7).

OROV assembly at the Golgi complex is stimulated by the cellular Endosomal Sorting Complexes Required for Transport (ESCRT) machinery (6). The ESCRT machinery is most commonly associated with membrane deformation and scission in an ‘inside-out’ topology, in contrast to the ‘outside-in’ topology of endocytic vesicles (9). The human ESCRT machinery comprises multiple distinct protein complexes (ESCRT-0, ESCRT-I, ESCRT-II and ESCRT-III) plus several accessory proteins, with final membrane scission being promoted by the concerted action of ESCRT-III and the ATPase VPS4 (10). The ESCRT-III machinery is formed by members of the charged multivesicular body protein (CHMP) family, with distinct CHMP proteins playing specific coordinated roles. The first ESCRT-III component recruited is CHMP6, which recruits CHMP4 isotypes (CHMP4A or CHMP4B) to homo-oligomerizes at the target membrane. The CHMP2A plus CHMP3 complex and/or CHMP2B alone are then recruited, activating VPS4 to promote ESCRT-III filament disassembly and membrane scission (10–12). Virus recruitment of the ESCRT-III machinery is canonically mediated by direct association of viral ‘late domains’ with ESCRT-I, the cellular ubiquitylation machinery and/or the accessory protein ALIX. These interactions culminate in recruitment and activation of the ESCRT-III machinery (13, 14). With the exception of flaviviruses, which recruit ESCRT components to the ER (15), most viruses recruit ESCRT machinery to the plasma membrane, endosomes or other ‘post-Golgi’ membranes (14, 16). It is therefore particularly interesting that OROV budding involves direct recruitment of ESCRT machinery to Golgi membranes (6). The OROV glycoproteins do not contain any identifiable viral ‘late domains’ and the molecular basis of ESCRT machinery recruitment by OROV thus remains to be established.

The study of many orthobunyaviruses requires high-level biosafety containment, owing to their ability to cause severe human disease and a lack of effective vaccines. Recombinant systems that recapitulate key aspects of virus biology are thus desirable to facilitate investigation under reduced biosafety containment of the assembly pathways of these highly-pathogenic virus (17, 18). Plasmid-based recombinant virus-like particle (VLP) production systems have been established for several members of the order *Bunyavirales*, with phlebovirus and hantavirus VLP assembly requiring expression of the viral glycoproteins but not the nucleoprotein (19, 20). Mini-replicon systems where the M polyprotein, L protein (RdRP) and nucleoprotein are recombinantly expressed on plasmids to generate VLPs has been reported for several orthobunyaviruses including BUNV, OROV, La Crosse virus (LACV), Akabane virus, and Schmallenberg virus (SBV) (21–27). Additionally, transcription- and replication-competent VLPs facilitate measurement of subtle differences in LACV and Rift Valley fever virus (RVFV) RdRp activity (27). Given that OROV recombinant virus lacking NSm is fully infectious (28) and that the signal peptide of NSm (SP^NSm^) is cleaved from the BUNV Gn protein during virion maturation (29), we hypothesised that recombinantly expressed OROV glycoproteins Gn and Gc would self-associate and be release from cells in a VLP-like manner.

Here we report OROV glycoprotein secretion via recombinant expression of the M polyprotein. The mature OROV Gn protein appears to contain two transmembrane domains (TMDs), both when expressed recombinantly and in the context of infection, and the second TMD is crucial for glycoprotein delivery to the Golgi and subsequent secretion. Pharmacological disruption of Golgi function prevents OROV glycoprotein secretion, as does perturbation of ESCRT function, highlighting the utility of this secretion assay for probing cellular determinants of OROV assembly. Finally, we show that the ESCRT-III component CHMP6 interacts with the OROV glycoproteins, suggesting that interaction between OROV glycoproteins and CHMP6 may facilitate OROV assembly and release.

## Results

### Processing and secretion of recombinant OROV glycoproteins

A synthetic codon-optimised cDNA encoding the OROV medium (M) segment polyprotein (residues 1– 1420) was cloned into the mammalian expression vector pcDNA3.1 with an HA epitope tag between the secretion signal sequence (SS; residues 1–16) and the first residue of the mature Gn protein (Fig. 1A). Transfection of HEK293T cells with a plasmid encoding HA-tagged OROV M polyprotein yielded high levels of Gn and Gc (observed as bands at ∼35 kDa and ∼120 kDa, respectively) in the supernatant, confirming secretion of OROV glycoproteins by these transfected cells (Fig. 1B). In order to establish whether both OROV glycoproteins are required for secretion, OROV Gn (M polyprotein residues 17–313) and Gc (M polyprotein residues 482–1420) were cloned separately into pcDNA3.1 after the OROV M SS and HA epitope tag (Fig. 1A), the polyprotein cleavage sites being inferred from homology with those determined or predicted for the BUNV M polyprotein (29). Domain boundaries were chosen based on prior studies of OROV and BUNV glycoproteins (29, 30) and analysis of predicted TMD topology (31). As expected, neither Gn nor Gc was abundant in the cell supernatant when transfected into HEK293T cells in isolation. However, we also failed to observe secreted glycoproteins when the proteins were co-expressed (Fig. 1B). This was unexpected as a previous study of OROV infection had shown that the NSm polypeptide, which lies between Gn and Gc in the M polyprotein, was dispensable for virion production (28). Furthermore, we observed that Gn from cells transfected with a plasmid encoding HA-tagged M polyprotein migrated more slowly in SDS-PAGE than Gn from cells transfected with the HA-Gn^313^ plasmid (Fig. 1B), consistent with the two proteins having different masses.

**Figure 1.**
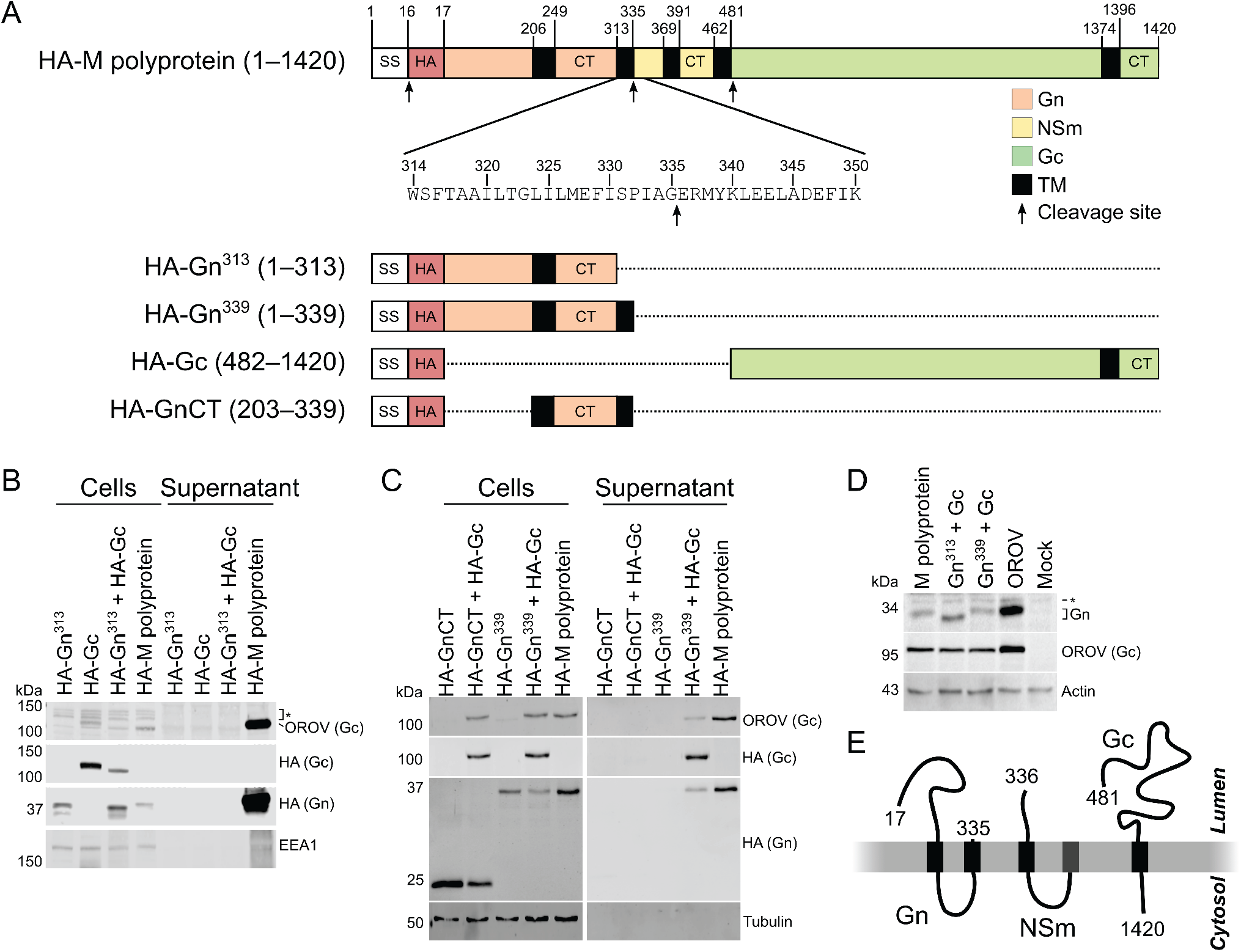
Recombinant production and secretion of OROV glycoproteins. (**A**) Schematic representation of the recombinant constructs used to express the OROV M polyprotein, individual glycoproteins or domains thereof. Expression constructs were preceded by the OROV M secretion signal sequence (SS; unshaded) and an HA epitope tag (HA; red shading). Regions corresponding to Gn, NSm and Gc are shaded in orange, yellow and green, respectively, with cytosolic tails denoted (CT). Transmembrane regions (black) as predicted by TMHMM (31) and polyprotein cleavage sites (arrows) are shown. (**B** and **C**) HEK293T cells were transfected with HA-tagged OROV M (poly)protein constructs. After 48 h cell lysates and supernatants were harvested, subjected to SDS-PAGE and immunoblotted using anti-HA and anti-OROV antibodies to detect secreted proteins, the latter polyclonal antibody detecting Gc but not Gn, and with anti-EEA1 (**B**) or anti-tubulin (**C**) antibodies used as loading controls. Data is representative of two (**B**) or three (**C**) independent experiments. Non-specific bands are marked (*). (**D**) HEK293T cells were transfected with OROV M (poly)proteins lacking an N-terminal epitope tag or infected with OROV (MOI = 1). After 24 h cells were harvested and subjected to SDS-PAGE and immunoblotting using antibodies shown, anti-actin serving as a loading control. Non-specific band is marked (*). Data is representative of four independent experiments. (**E**) Schematic representation of OROV M polyprotein processing.

BUNV Gn maturation occurs in a stepwise fashion, with Gn first being liberated from the rest of the M polyprotein via cleavage by signal peptidase between residues 331–332 (equivalent to OROV residues 335–336) to produce an immature Gn protein with two predicted TMDs, the second of TMD representing the NSm signal peptide (SP^NSm^) and being subsequently removed by signal peptide peptidase (SPP) to yield mature Gn with a carboxy terminus at approximately residue 312 (29). Analysis of the OROV M polypeptide sequence using the SignalP 5.0 server (32) suggested that OROV Gn may also be liberated from the M polyprotein via cleavage by signal peptidase between residues G335 and E336 (Fig. 1A, second arrow). We therefore exploited an additional expression construct we had generated that included this predicted TMD, comprising the OROV M SS and HA-tag followed by M polypeptide residues 17–339 (Gn^339^). While Gn^339^ was not secreted into the medium following expressed on its own in HEK293T cells, both Gn^339^ and Gc were abundant in the supernatant of cells co-transfected with constructs encoding both proteins (Fig. 1C). Co-expression of Gc with Gn^339^ is thus sufficient to promote release of both proteins from cells, presumably as VLPs or as exocytic vesicles. To test whether the two predicted TMDs plus cytosolic region of Gn^339^ were sufficient to promote Gc secretion, HEK293T cells were co-transfected with expression constructs encoding HA-tagged Gc and Gn residues 203– 339 (GnCT). Neither Gc nor GnCT was secreted to the supernatant of these co-transfected cells, indicating that GnCT is not sufficient to promote secretion of Gc (Fig. 1C). To allow direct comparison between Gn produced via recombinant expression and following OROV infection, an antibody was raised against Gn residues 99–112 and recombinant constructs encoding Gc, Gn^313^, Gn^339^ and the M polyprotein with the OROV SS but no N-terminal epitope tag were cloned. Immunoblotting of HEK293T cells transfected with M polyprotein, Gn^313^+Gc, Gn^339^+Gc, or infected with OROV (MOI = 1) demonstrates that Gn from cells transfected with Gn^339^, with M polyprotein or infected with OROV all display equivalent electrophoretic mobility, whereas Gn^313^ migrates substantially faster (Fig. 1D). Together, these results demonstrate that recombinant OROV glycoproteins are secreted when co-expressed as individual proteins or as a polyprotein, but unlike BUNV the second TMD of OROV Gn is not liberated by SPP following either recombinant expression or infection (Fig 1E).

### The Gn^339^:Gc complex is sufficient for Golgi localization

During infection, the orthobunyavirus M polyprotein is translated at the ER before the Gn and Gc proteins are transported to the Golgi apparatus, which is the main site of virus assembly (5). In order to investigate the subcellular distribution of recombinant OROV glycoproteins (Fig. 1A), HeLa cells were transfected with HA-tagged OROV glycoproteins and then stained for the HA signal and for the ER chaperone calnexin 2 (CNX2) or the *cis*-Golgi marker GM130 (Fig. 2). When expressed alone, HA-tagged Gn^313^, Gc and Gn^339^ all co-localized with CNX2 at the ER and did not co-localise with GM130 at the Golgi (Fig. 2A–C). Similarly, when HA-tagged Gn^313^ and Gc were co-expressed they were retained at the ER and did not co-localise at the Golgi (Fig. 2D). However, when HA-tagged Gn^339^ and Gc were co-expressed they showed extensive co-localisation with GM130 at the Golgi and less-extensive co-localisation with CNX2 at the ER (Fig. 2E). Extensive GM130 co-localisation was also observed when the HA-tagged OROV M polyprotein was expressed (Fig. 2F). Additionally, HA-stained puncta could often be observed outside cells transfected with M polyprotein (Fig. 2F), suggesting that secreted glycoproteins adhered to the coverslip during fixation and subsequent preparation of the microscope slides. Taken together, these data indicate that co-expression of Gc with Gn spanning the first two predicted TMDs (residues 1–339; Gn^339^), either as separate polypeptides or as a polyprotein, is necessary and sufficient for these proteins to be transported from the ER to the Golgi apparatus.

**Figure 2.**
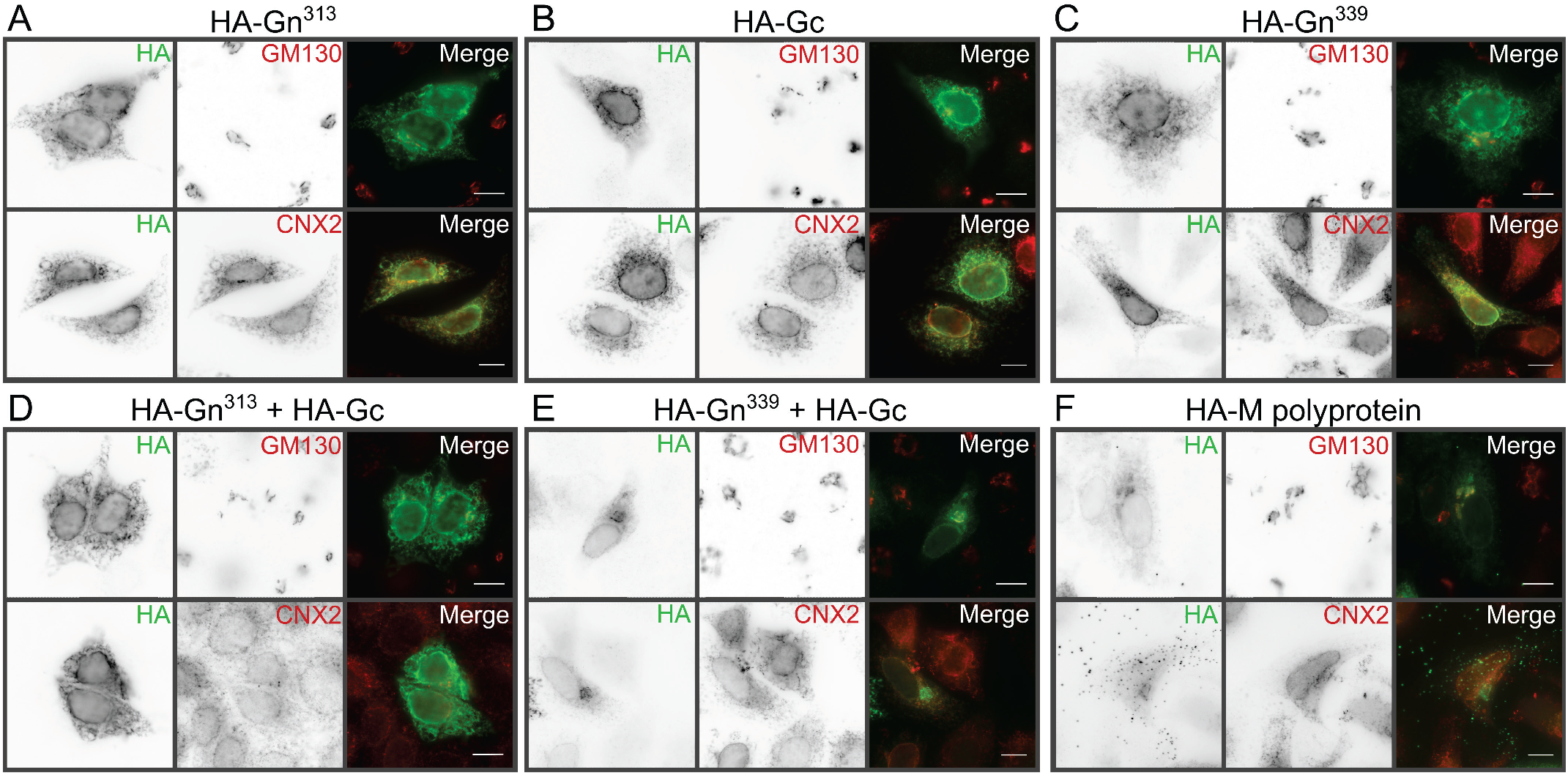
Intracellular localization of OROV glycoproteins. (**A** to **F**) HeLa cells were transfected with plasmids expressing the HA-tagged OROV glycoproteins or the M polyprotein (as indicated). Cells were fixed and co-stained with rabbit anti-HA (AF488) and either mouse anti-GM130 (AF568) or mouse anti-CNX2 (AF568) antibodies to identify Golgi and ER compartments, respectively, before analysis using wide-field fluorescence microscopy. Scale bars = 10 µm.

### Inhibition of the post-Golgi transport disrupts OROV glycoprotein secretion

The OROV assembly pathway involves the recruitment of the host ESCRT machinery to Golgi membranes for virus particle production (6). To further investigate the intracellular trafficking of the OROV M polyprotein, two drugs were used to interfere with different steps of Golgi transport: Brefeldin A was used to inhibit ER to Golgi transport, and monensin was used to block post-Golgi transport (33, 34). HEK293T cells were transfected with HA-tagged OROV M and the culture medium was collected before and after drug treatment to monitor glycoprotein secretion. Secretion of OROV glycoproteins was severely disrupted in cells treated with monensin or brefeldin A (Fig. 3A,B). Microscopic analysis of drug-treated HeLa cells using the *cis*-Golgi marker GM130 (Fig. 3C) showed that brefeldin A caused the anticipated dispersal of the *cis*-Golgi (35) and it prevented co-localisation of GM130 with the OROV glycoproteins. Conversely, monensin treatment led to strong co-localisation of the glycoproteins with enlarged and/or fragmented GM130-positive structures (Fig. 3C), osmotic swelling and fragmentation of the Golgi being a consequence of monensin treatment (34). These results demonstrate that correct Golgi function is required for OROV glycoprotein secretion.

**Figure 3.**
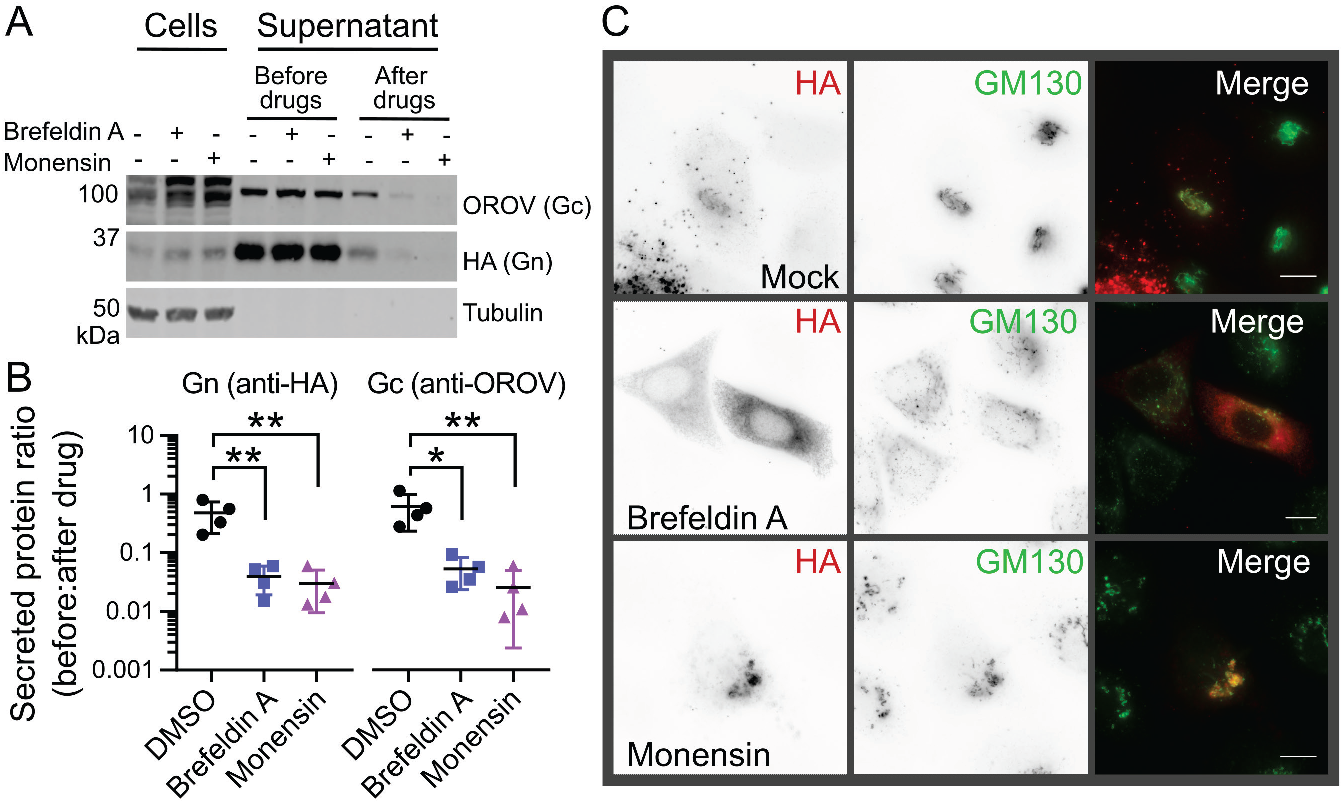
Disruption of the Golgi apparatus prevents OROV glycoprotein secretion. (**A**) HEK293T cells were transfected with HA-tagged OROV M polyprotein and after 18 h the cell supernatant was harvested and reserved. The culture medium was replenished and supplemented with 5 µg/mL brefeldin A, 1 µM monensin or DMSO as a control. After 6 h, cells and supernatant were harvested. All samples were then subjected to SDS-PAGE and immunoblotted using antibodies shown, anti-tubulin acting as a loading control. (**B**) Ratio of Gn (anti-HA) and Gc (anti-OROV) protein present in supernatants collected before or after drug treatment. Mean ± SD is shown for four independent experiments. *, p ≤ 0.05; **, p ≤ 0.01 (One-way ANOVA using Dunnett’s test for multiple comparisons to DMSO control cells). (**C**) HeLa cells transfected and drug treated as in (**A**) before fixation, co-staining with rabbit anti-HA (AF568) plus mouse anti-GM130 (AF488) to identify Golgi membranes and analysis using wide-field fluorescence microscopy. Scale bars = 10 µm.

### OROV glycoproteins secretion requires ESCRT machinery

During OROV infection, multiple ESCRT components are recruited to Golgi membranes in order to promote virus particle production (6). VPS4 is a cellular ATPase that stimulates the disassembly of assembled ESCRT-III filaments at the necks of vesicles budding away from the cytoplasm, promoting membrane scission (10). VPS4 is recruited to Golgi membranes during OROV infection and the morphology of OROV viral factories is altered when infected cells express a dominant negative form of this enzyme (VPS4E/Q) that inhibits ESCRT-III disassembly (6). In order to analyse whether secretion of OROV glycoproteins is VPS4-dependent, HA-tagged OROV M polyprotein was transfected into HEK293 cells stably expressing GFP-tagged wild-type human VPS4 or a dominant negative E228Q mutant (VPS4E/Q) under the control of ecdysone response elements (36). Following transfection with plasmids encoding HA-tagged OROV M polyprotein these VPS4-expressing cells, and the parental control cells, were treated with either ponasterone A (Pon A) to induce VPS4 expression or with DMSO as a vehicle control and the abundance of cell-associated and secreted OROV proteins was monitored by immunoblotting (Fig. 4A). The secretion of Gn and Gc by parental cells and those expressing wild-type GFP-VPS4 did not significantly differ when cells were treated with Pon A or DMSO (Fig. 4B). Gn and Gc secretion was decreased when GFP-VPS4E/Q cells were treated with Pon A, although only the decrease in Gn was statistically significant (Fig. 4B). To probe whether VPS4 activity was also required for OROV virus particle release, the VPS4-expressing HEK293 cells were infected with OROV (MOI = 1) for 4 h before induction of VPS4 expression and analysis of cell-associated plus secreted OROV protein abundance at 24 h post-infection (Fig. 4C). Both Gc and N protein secretion was significantly decreased in GFP-VPS4E/Q cells treated with Pon A following infection with OROV, whereas Pon A treatment of the parental or GFP-VPS4wt expressing cells did not alter OROV protein secretion (Fig. 4D).

**Figure 4.**
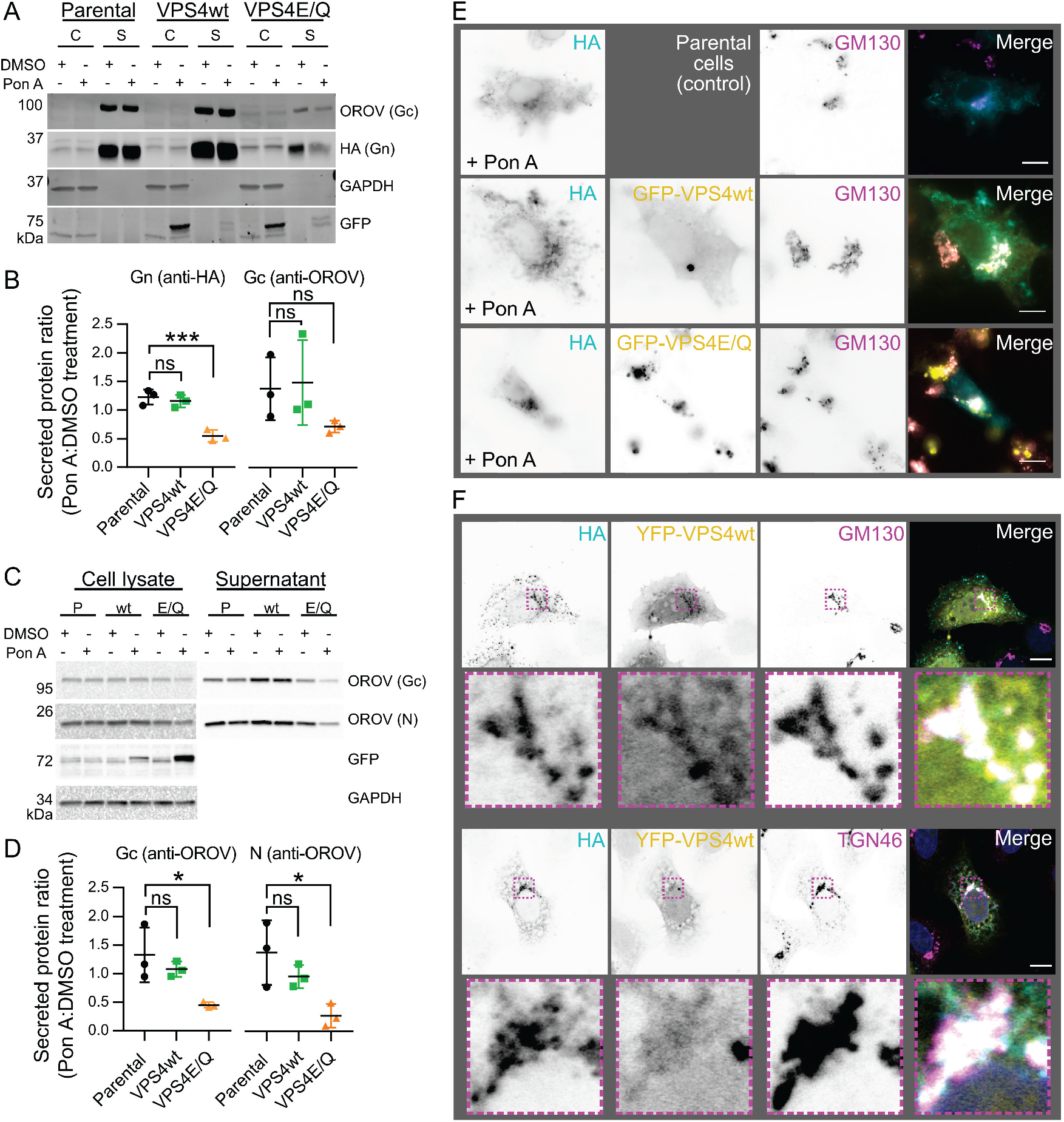
OROV protein secretion depends on VPS4 activity. (**A**) HEK293 cells stably expressing GFP-tagged wild-type VPS4 (VPS4wt) or a dominant negative mutant (VPS4E/Q) under the control of the ecdysone response element, or parental cells as a control, were transfected with HA-tagged M polyprotein and treated with either 1 µM ponasterone A (Pon A) or with DMSO as a control. After 48 h, cells (C) and supernatant (S) were harvested, subjected to SDS-PAGE and immunoblotted using antibodies shown. (**B**) Quantitation of the ratio of OROV Gn (anti-HA; left) or Gc (anti-OROV; right) that was secreted following treatment with Pon A versus DMSO. Mean ± SD from three independent experiments is shown. ***, p ≤ 0.001 (One-way ANOVA using Dunnett’s test for multiple comparisons to parental control cells). (**C**) HEK293 cells stably expressing VPS4wt (wt) or VPS4E/Q (E/Q) under the control of the ecdysone response element, or parental cells (P) as a control, were infected with OROV (MOI = 1) and after 4 h were treated with 1 µM Pon A or DMSO as a control. After 24 h cells and supernatant were harvested, subjected to SDS-PAGE and immunoblotted using antibodies shown, anti-OROV detecting both N and Gc proteins. (**D**) Quantitation of the ratio of OROV Gc (left) and N (right) secretion in cells treated with Pon A versus DMSO. Mean ± SD from three independent experiments is shown. *, p ≤ 0.05 (One-way ANOVA using Dunnett’s test for multiple comparisons to parental control cells). (**E**) HEK293 cells expressing GFP-tagged VPS4wt or VPS4E/Q, or parental cells, were transfected with HA-tagged OROV M polyprotein and treated with 1 µM Pon A for 18 h before being fixed and co-stained with rabbit anti-HA (AF568) and mouse anti-GM130 (AF647) antibodies and then analysed by wide-field fluorescence microscopy. Scale bars = 10 µm. (**F**) HeLa cells were co-transfected with plasmids expressing HA-tagged OROV M polyprotein and YFP-VPS4wt. After 24 h cells were fixed and co-stained with rabbit anti-HA (AF568) and (top) mouse anti-GM130 (AF647) or (bottom) sheep anti-TGN46 (AF647), and then analysed by confocal microscopy. Scale bars = 10 µm.

OROV glycoproteins and GFP-tagged VPS4E/Q co-localised near GM130-positive *cis*-Golgi membranes when the Pon A treated HEK293 stable cells were transfected with HA-tagged OROV M polyprotein (Fig. 4E), similar to previous observations in OROV-infected HeLa cells (6), but unlike previous studies GFP-tagged wild-type VPS4 did not co-localise with either OROV glycoproteins or the Golgi marker GM130 (Fig. 4E). However, there was partial co-localisation between OROV glycoproteins and wild-type VPS4 near Golgi (GM130- and TGN46-positive) membranes when HeLa cells were co-transfected with YFP-tagged VPS4 and OROV M polyprotein (Fig. 4F). Taken together, these results are consistent with VPS4 activity being required for OROV glycoprotein/virion secretion and with the extent of fluorescently-tagged VPS4 localisation at the Golgi differing between HEK293 and HeLa cells.

To further investigate the interaction between ESCRT-III components and OROV glycoproteins, HeLa cells were transiently co-transfected with HA-tagged OROV M polyprotein and dominant-negative C-terminally YFP tagged members of the charged multivesicular body protein (CHMP) family (37). Although no co-localisation could be detected with the majority of CHMP proteins, partial co-localisation could be observed between OROV glycoproteins and CHMP6-YFP (Fig. 5). A similar pattern of partial co-localisation could be observed between OROV glycoproteins and endogenous CHMP6 (Fig. 6A), the co-localisation of both endogenous and YFP-tagged CHMP6 with OROV glycoproteins being more prominent when cells were treated with monensin to stall glycoprotein trafficking at the Golgi (Fig. 6A,B).

**Figure 5.**
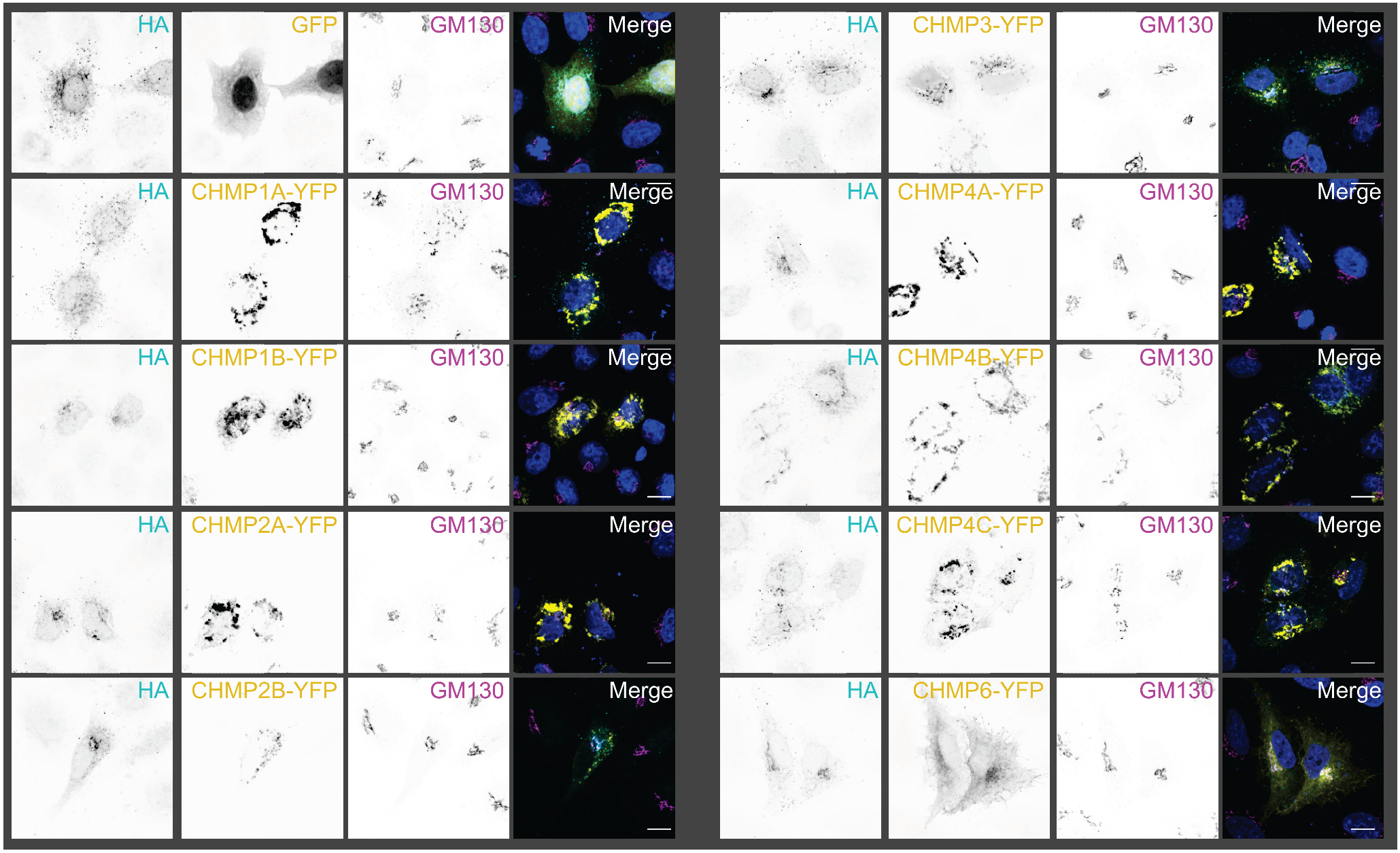
OROV glycoproteins co-localise with CHMP6 but not with other ESCRT-III components. HeLa cells were co-transfected with plasmids expressing HA-tagged OROV M polyprotein and YFP-tagged CHMP proteins (as indicated). After 24 h cells were fixed and co-stained with rabbit anti-HA (AF568) and mouse anti-GM130 (AF647), and analysed by confocal microscopy. Scale bars = 10 µm.

**Figure 6.**
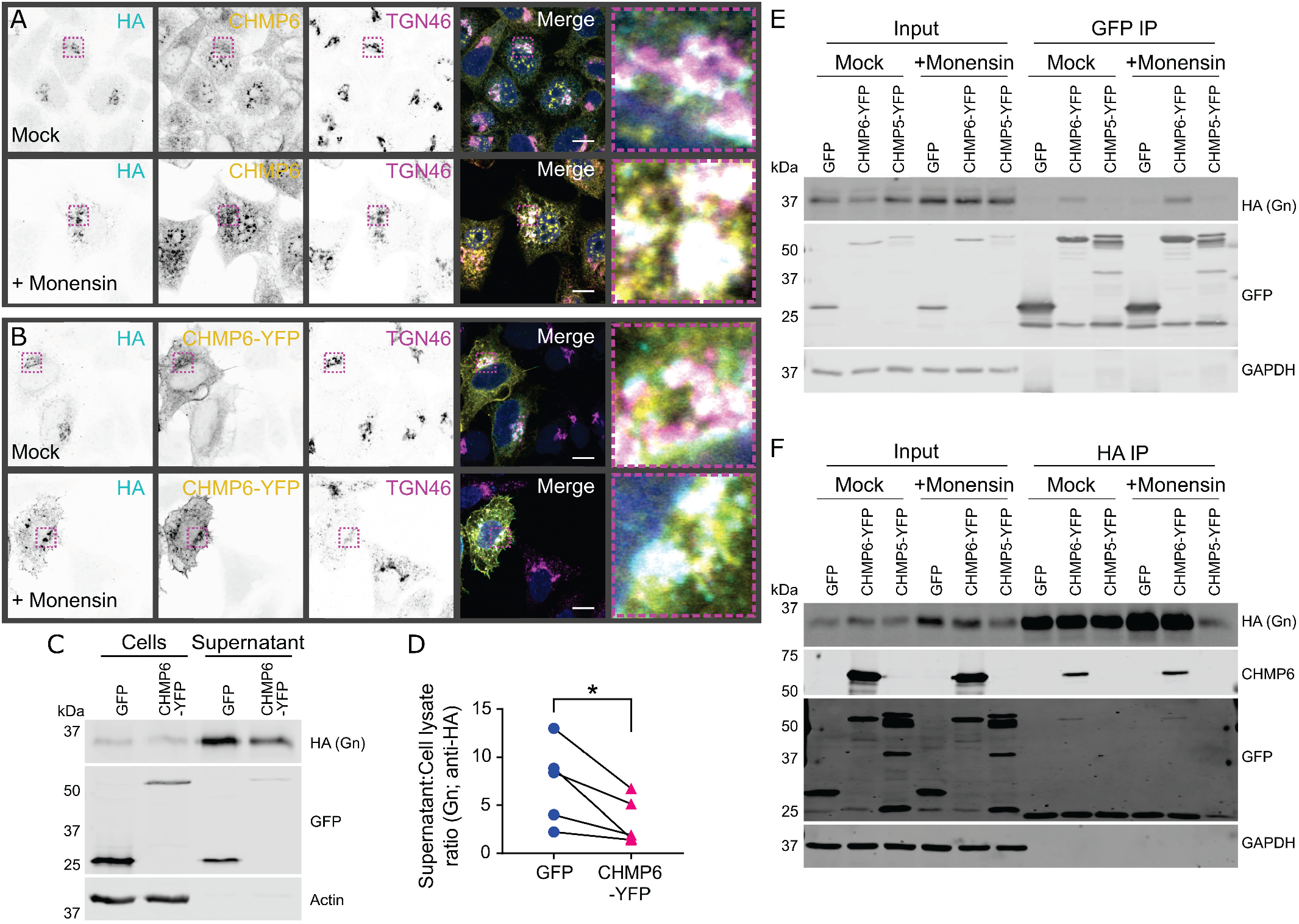
OROV glycoproteins interact with CHMP6. (**A**) HeLa cells were transfected with HA-tagged OROV M polyprotein. After 18 h, cells were treated with 1 µM monensin or mock-treated for 6 h before being fixed and co-stained with mouse anti-HA (AF568), rabbit anti-CHMP6 (AF488) and sheep anti-TGN46 (AF647) before analysis by confocal microscopy. Scale bars = 10 µm. (**B**) HeLa cells were co-transfected with HA-tagged OROV M polyprotein and YFP-tagged CHMP6. Cells were drug treated and fixed as in (A) before staining with rabbit anti-HA (AF568) and sheep anti-TGN46 (AF647) before analysis by confocal microscopy. Scale bars = 10 µm. (**C**) HEK293T cells were co-transfected with HA-tagged M polyprotein and GFP or CHMP6-YFP. After 48 h cells and supernatant were harvested, subjected to SDS-PAGE and immunoblotted using antibodies listed. (**D**) Ratio of Gn (anti-HA) signal in the culture supernatant versus cell lysate. Data are from five independent experiments. *, p ≤ 0.05 (paired ratio t test). (**E**,**F**) HEK293T cells were co-transfected with HA-M polyprotein and either GFP, CHMP6-YFP or CHMP5-YFP. After 18 h, cells were treated with 1 µM monensin or mock-treated for 6 h. Cells were harvest and lysates subjected to affinity capture (IP) using either GFP (**E**) or HA (**F**) affinity matrices before SDS-PAGE and immunoblotting using the antibodies shown. Representative blots from two (**E**) or three (**F**) independent experiments are shown.

CHMP6 nucleates the polymerisation of ESCRT-III filaments (9) and overexpression of CHMP6-YFP confers a mild reduction in ESCRT-mediated secretion of HSV-1 and HIV-1 (38). OROV glycoprotein secretion is significantly impaired when HEK293T cells are co-transfected with HA-tagged M polyprotein and CHMP6-YFP versus GFP (Fig. 6C,D). Immunoprecipitation of co-transfected HEK293T cells demonstrates that CHMP6 physically interacts with OROV glycoproteins: HA-tagged OROV glycoproteins co-precipitate with CHMP6-YFP captured on GFP affinity resin (Fig. 6E) and CHMP6-YFP co-precipitates with OROV glycoproteins captured using HA affinity resin (Fig. 6F). In both cases co-precipitation is not observed with GFP or with CHMP5-YFP, confirming that the interaction is specific. These data demonstrate that the ESCRT-III component CHMP6 interacts with OROV glycoproteins, and that this interaction may promote glycoprotein secretion.

## Discussion

We have characterized the maturation and secretion of OROV glycoproteins to study virus assembly and budding. Surface glycoproteins are the principal viral components necessary for VLP production across other members of the order *Bunyavirales* (19, 20). We demonstrate that OROV glycoproteins are secreted when Gc is co-expressed with Gn but that the proteolytic processing of OROV Gn differs from that of other orthobunyaviruses. Glycoprotein secretion is inhibited by drugs that interfere with Golgi function. Secretion is also inhibited when the cellular ESCRT machinery is disrupted, and the ESCRT-III component CHMP6 physically interacts with OROV glycoproteins.

Previous studies of BUNV have shown two cleavage sites between the boundaries of Gn and NSm in the M polyprotein, either side of the second TMD (29). Specifically, Shi and colleagues (29) observed that the second TMD of the BUNV M polyprotein (which they term SP^NSm^) acts as a signal peptide for NSm, being cleaved at its C terminus by cellular signal peptidases, and that this TMD is subsequently liberated from the mature Gn protein by SPP. The migration of BUNV Gn in SDS-PAGE changed linearly when Gn was truncated before the start of the second TMD, but the recombinant protein migrated identically to mature Gn from BUNV infection or M polyprotein expression when stop codons were introduced at or following the start of this second TMD. In contrast, we observe that the electrophoretic mobility of Gn^339^, which includes the second predicted TMD, matches that of Gn from M polyprotein expression or OROV infection whereas the migration of untagged OROV Gn^313^ is faster, consistent with Gn^313^ being a truncated form of Gn (Fig. 1D). Furthermore, we observe the efficient secretion of OROV glycoproteins into the culture medium when Gn^339^ is co-expressed with Gc or when OROV M is expressed as a polyprotein, but not when Gn^313^ is co-expressed with Gc. Taken together, these data show that the OROV M polyprotein is processed to a mature Gn protein spanning two TMDs (Gn^335^), both in the context of recombinant polyprotein expression and during OROV infection. This contrasts with previous observations for BUNV and suggests that the individual orthobunyaviruses have differing requirements for cleavage of the second Gn TMD by SPP.

In addition to differences in proteolytic processing of the M polyprotein between BUNV and OROV, we observed a difference in the requirement for co-expression of Gn and Gc in order for these proteins to traffic from the ER to the Golgi. BUNV Gn is trafficked to the Golgi complex when expressed alone (8) due to the Golgi retention signal located on its first TMD (39). In contrast, we observe that neither Gn with either one or two predicted TMDs (Gn^313^ and Gn^339^, respectively) nor Gc localise to the Golgi when expressed in isolation. Co-expression of Gc with Gn^339^ (two predicted TMDs) resulted in strong co-localisation of the glycoproteins with the *cis*-Golgi marker GM130, as did expression of the entire M polyprotein. The difference in requirement for Gn and Gc co-expression in order to mature beyond the ER into the Golgi may highlight structural differences between the orthobunyavirus glycoproteins, with the BUNV Gn extracellular domain being capable of folding independently whereas the OROV Gn extracellular region requires Gc for folding and/or stabilisation.

Since bunyaviruses do not possess matrix proteins, the cytosolic tails of Gn and Gc are believed to be important for virus assembly and budding. The Gn cytosolic domain of Uukuniemi phlebovirus interacts with virus ribonucleoprotein, which is crucial for genome packaging (40). Furthermore, the Gn and Gc cytosolic tails of BUNV are required for Golgi complex targeting, virus assembly and infectivity (41), with the first TMD of BUNV Gn being required for correct Golgi targeting of Gc (39). We therefore investigated whether the Gn cytosolic tail (residues 203–339, GnCT) could serve as a chaperone for Gc. Gc is not secreted into the medium when co-expressed with GnCT, whereas co-expression of Gc with Gn^339^ did promote glycoprotein secretion (Fig. 1C). This data confirms that the Gn^339^ extracellular domain is required for secretion of Gc, consistent with previous studies of BUNV showing that mutations in the glycosylation sites of Gn prevented correct folding of both Gn and Gc (42).

Previous studies of BUNV showed that budding and initial maturation steps occur in the Golgi stacks, but that full virus maturation does not proceed without a functional *trans*-Golgi (7). Fragmentation of the Golgi is observed when Vero cells are infected with BUNV, although these fragmented Golgi are still competent to sustain infectious virion production (7). Similarly, OROV infection leads to *trans-*Golgi network fragmentation as detected by dispersion of the TGN46 marker (6). We do not observe fragmentation of the Golgi in cells expressing the M polyprotein (Figs 2F and 6A), suggesting that additional OROV proteins are required to promote Golgi fragmentation. Two drugs that interfere with the Golgi complex via different mechanisms were used to investigate the intracellular trafficking of OROV glycoproteins. The drug monensin strongly disrupts *trans*-Golgi network function by altering the pH (34), allowing us to probe post-Golgi transport of glycoproteins (Fig. 3). We observed an accumulation of intracellular glycoproteins at GM130-positive compartments in monensin-treated cells with a concomitant drop in glycoprotein secretion into the extracellular medium, confirming that a functional *trans*-Golgi network is required for OROV glycoprotein maturation and secretion. Brefeldin A prevents protein traffic from the ER to the Golgi but allows retrograde trafficking to proceed, causing re-adsorption of the Golgi into the ER (35). As expected, we observed a loss of the distinct Golgi-like staining patterns for both the OROV glycoproteins and the *cis*-Golgi marker GM130, plus a loss of glycoprotein secretion, following brefeldin A treatment (Fig. 3C). Interestingly, we do not observe accumulation of intracellular glycoproteins, consistent with a loss in protein translation caused by the ER stress induced by brefeldin A treatment (43). Taken together, our data show disruption of Golgi function inhibits OROV glycoprotein maturation and secretion, as has been observed previously for cells infected with BUNV (44).

The production of OROV infectious particles requires the cellular ESCRT machinery (6). To probe whether OROV glycoprotein secretion also requires ESCRT activity, and thus faithfully recapitulates infectious virus particle production, we investigated the association of ESCRT components with OROV glycoproteins and role of the cellular ESCRT machinery in viral glycoprotein secretion. The ESCRT-III– disassembling ATPase VPS4 co-localises with OROV replication sites, expression of the ATPase-defective VPS4E/Q mutant leading to changes in replication compartment morphology (6). Secretion of OROV glycoproteins is decreased when cells express VPS4E/Q, whereas secretion from parental cells or those expressing wild-type VPS4 is unchanged (Fig. 4A,B). Furthermore, a similar decrease in secretion of OROV Gc and N (presumably as virions) is observed during infection in VPS4E/Q-expressing cells versus parental and wild-type VPS4 expressing cells (Fig. 4C,D). This strongly suggests that VPS4 activity promotes OROV viral glycoprotein and virion secretion. Interestingly, while GFP-VPS4E/Q co-localized with OROV glycoproteins at Golgi membranes in HEK293 cells (Fig. 4E) in a similar manner to OROV infectious particles (6), GFP-VPS4wt did not. However, overexpression in HeLa cells yields co-localisation of YFP-VPS4wt with OROV glycoproteins and Golgi membranes (Fig. 4F), as observed previously for OROV infectious particles (6), consistent with the steady-state abundance of co-localised OROV glycoproteins and VPS4 differing between different cell types.

To further understand how cellular ESCRT components may promote OROV glycoprotein secretion, we investigated the co-localisation of OROV glycoproteins with overexpressed, dominant-negative YFP-tagged cellular ESCRT-III components (Fig. 5). Partial co-localisation was observed with both YFP-tagged and endogenous CHMP6 (Figs 5 and 6A), a myristoylated protein that nucleates ESCRT-III polymerisation (9), and co-immunoprecipitation experiments demonstrate a physical interaction between OROV glycoproteins and CHMP6-YFP (Fig. 6E,F). Most enveloped viruses that recruit the cellular ESCRT membrane-remodelling machinery for budding do so using peptide motifs known as viral ‘late domains’ (13, 14), but the OROV structural proteins lack identifiable late domain sequences. The Bro1-domain containing protein ALIX and ESCRT-I component TSG101, which act in parallel to stimulate ESCRT-III polymerisation (9), both contribute to OROV viral factory morphogenesis and virus production (6). Demonstration that OROV glycoproteins physically interact with CHMP6 raises the interesting hypothesis that OROV might also recruit CHMP6 directly to further enhance ESCRT-III polymer formation on Golgi membranes. Hijacking of cellular ESCRT activity via by direct stimulation of ESCRT-III polymerisation, rather than via late domain mediated recruitment of ESCRT adaptors, has been hypothesised for herpesviruses (45, 46) but remains to be shown definitively. Overexpression of CHMP6-YFP significantly decreases the secretion of OROV glycoproteins (Fig. 6C,D). While this is consistent with the tagged CHMP6 protein having a dominant negative effect upon ESCRT-mediated OROV glycoprotein secretion, it does not differentiate between ESCRT-III activity being stimulated via ESCRT-I/ESCRT-II mediated recruitment or by direct CHMP6 binding to OROV glycoproteins. Further experiments are required to dissect the molecular mechanisms by which ESCRT-III activity is stimulated to promote OROV glycoprotein and virion secretion.

In summary, we have demonstrated that OROV Gn contains two predicted TMDs that are essential for OROV glycoprotein trafficking to the Golgi complex. Pharmacological disruption of Golgi function inhibits OROV glycoprotein secretion, as do perturbations of ESCRT complex function. Furthermore, we observe co-localisation and interaction of CHMP6 of the ESCRT-III components with the OROV glycoproteins, suggesting that OROV may directly stimulate ESCRT-III polymerisation to support virus budding. *In vitro* expression of OROV glycoproteins represents a convenient experimental system for analysing the cellular determinants of OROV assembly and budding, avoiding requirements for high-containment biosecurity. The similarity in glycoprotein processing and requirement for VPS4 activity suggests that OROV Gn and Gc co-expression outside the context of infection leads to glycoprotein self-assembly and secretion from the cell as VLPs, although formal verification of this awaits future electron microscopy analysis.

## Materials and Methods

### Cell Culture

Mycoplasma-free HeLa and HEK293T cells were maintained in Dulbeco’s modified Eagle’s medium (DMEM; Gibco) supplemented with 10% heat-inactivated foetal calf serum (FCS) and 2 mM glutamine at 37°C in a humidified 5% CO2 atmosphere. HEK293 cells stably expressing the ecdysone receptor (EcR-293; Invitrogen), and derived cell lines stably expressing GFP-tagged wild-type or dominant negative (E228Q) human VPS4 under the control of ecdysone response elements, were maintained in the medium listed above supplemented with 400 µg/ml Zeocin and 800 µg/ml G418 (36).

### Plasmids and Constructs

All fragments of the M polyprotein from OROV strain BeAn19991 (GenBank ID KP052851.1) used in this study were synthesised (GeneArt) following codon-optimization for expression in human cells. Constructs encoding Gn^313^ (M polyprotein residues 1–313) and Gc (residues 482–1420) were synthesised with an HA epitope tag and BamHI restriction sequence immediately following the Gn secretion signal sequence (SS), and were cloned into pcDNA3.1(+) using the restriction enzymes NheI and XhoI. To generate the HA-tagged OROV M polyprotein construct, an extended NSm fragment (M polyprotein residues 299–496) was synthesised to overlap with the N and C termini of Gc and Gn^313^, respectively. The sequences for Gc, NSm and the entire pcDNA3.1 vector encoding Gn with an N-terminal SS plus HA tag were amplified by PCR using KOD Hot Start DNA polymerase (Novagen), mixed and then concatenated using the NEBuilder HiFi assembly kit according to the manufacturer’s instructions to yield pSnH-OROV-M. HA-tagged Gn^339^ was generated by site directed mutagenesis of pSnH-OROV-M to insert a stop codon after residue 339 using the mutagenic primers listed in Table 1. HA-tagged GnCT was amplified from pSnH-OROV-M using the primers shown in Table 1, digested with the restriction endonucleases BamHI and Xhoi, and ligated into BamHI/XhoI digested pSnH-OROV-M to yield the Gn SS and an HA epitope tag followed by M polyprotein residues 203–339. All expression constructs were verified by Sanger sequencing. CHMP-YFP plasmids were as described previously (38).

**Table 1.**
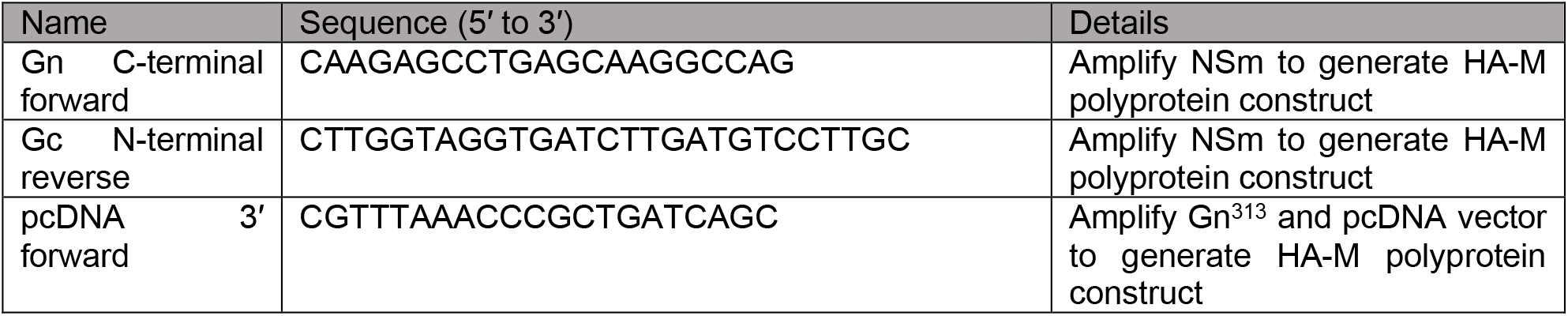
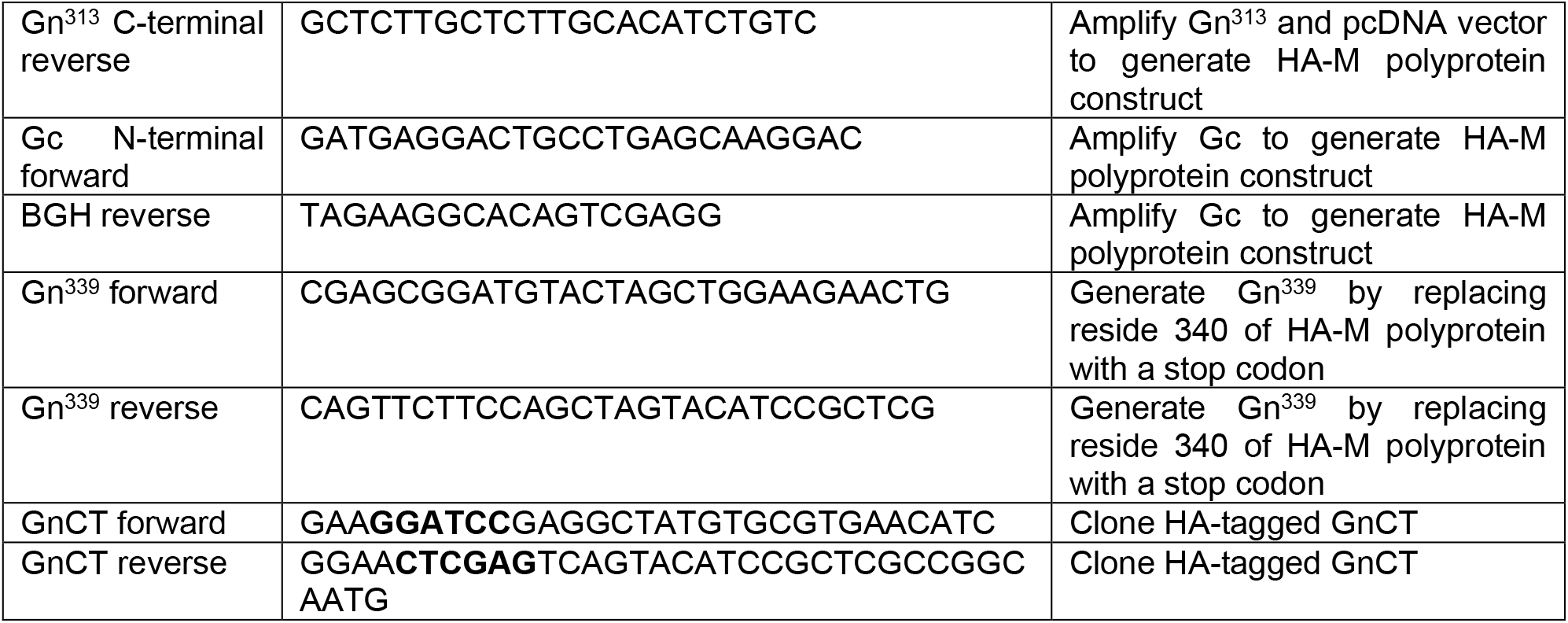
Oligonucleotide primers used in this study. Restriction sites are in **bold**.

### Antibodies

For immunoblot analyses the following primary antibodies (identifier, dilution) were used: anti-GFP (Sigma G2544, 1:5000), anti-OROV (Gc detection; ATCC VR-1228AF, 1:1000), anti-OROV antiserum (Gc and N detection; kindly donated by Luis Tadeu Moraes Figueiredo, University of São Paulo, 1:500), anti-CHMP6 (kindly donated by Wesley Sundquist, University of Utah, 1:1000), anti-HA (Cell Signaling Technology C29F4, 1:20,000), anti-actin (Sigma A3853, 1:1000), anti-tubulin hybridoma supernatant (clone YL1/2) (47), anti-GAPDH (GeneTex GTX28245, 1:10000) and anti-EEA1 (Abcam Ab2900, 1:1000). The secondary antibodies were LI-COR IRDye 680T conjugated donkey anti-rabbit (926-68023) or goat anti-mouse (926-68020), or LI-COR IRDye 800CW conjugated donkey anti-rabbit (926-32213) or goat anti-mouse (926-32210). Polyclonal serum against Gn was raised commercially (GenScript) by immunisation of rabbits with a synthetic peptide (Gn residues 99–112, sequence VLSVDDNGHIIPKMC) coupled to keyhole limpit hemocyanin, followed by affinity purification. This antibody was validated by immunoblotting lysates of HEK293T cells transfected with pSnH-OROV-M and was used at 1:1000 dilution. For immunocytochemistry the following primary antibodies (identifier, dilution) were used: anti-HA (Cell Signaling Technology C29F4, 1:1000), anti-GM130 (BD Biosciences 610822, 1:1000), anti-TGN46 (BioRad AHP500G, 1:250), anti-CHMP6 (kindly donated by Wesley Sundquist, University of Utah, 1:200), and anti-Calnexin2 (BD Biosciences 610523, 1:100). Secondary antibodies were Alexa Fluor conjugated goat anti-mouse (A-21236, A-11001 and A-11031, ThermoFisher), donkey anti-rabbit (A-10042, ThermoFisher), goat anti-rabbit (A-11008, ThermoFisher), donkey anti-mouse (A32766, ThermoFisher), and donkey anti-sheep (A21448, ThermoFisher).

### OROV glycoprotein secretion and immunoblotting

HEK293T cells were seeded at 2.5×10^5^ cells per well in six well dishes. Cells were transfected by mixing 2 µg of DNA (split evenly by mass between the plasmids indicated) and 1.5 μg of branched polyethylenimine (PEI; average MW ∼25,000, Merck) in Opti-MEM (ThermoFisher), incubating at room temperature for 20 min and then applying to cells. Cell culture medium was harvested after 48 h and cleared of cellular debris by centrifugation at 800×g for 10 min at 4°C. Glycoproteins in the supernatants were pelleted by centrifugation at 100,000×g at 4°C for 90 min using a TLA-55 rotor (Beckman). The resultant pellets, containing glycoproteins, were resuspended and boiled in sodium dodecyl sulfate-polyacrylamide gel electrophoresis (SDS-PAGE) sample buffer. Cell samples were harvested by centrifugation and resuspended in lysis buffer (10 mM Tris pH 7.5, 150 mM NaCl, 0.5% IGEPAL CA-630 [a.k.a. NP-40], 0.5 mM EDTA) supplemented with EDTA-free protease inhibitor cocktail (Roche) and incubated on ice for 30 min. Lysates were clarified by centrifugation at 20,000×g for 10 min at 4°C. The protein concentration in each lysate sample was normalized following quantification using the BCA assay (ThermoFisher) and samples were boiled in SDS-PAGE sample buffer. Samples were separated by SDS-PAGE using 10% or 15% (w/v) polyacrylamide gels and transferred to Protran nitrocellulose membranes (Perkin Elmer) using the Mini-Trans blot systems (BioRad) following the manufacturer’s protocol. After blocking in Intercept blocking buffer (LI-COR) for anti-Gn or PBS with 5% (w/v) non-fat milk powder for all other antibodies, membranes were incubated with primary antibody at room temperature for 1 h or overnight at 4°C, washed, and then incubated with the secondary antibody for 1 h at room temperature. Dried blots were visualized on an Odyssey CLx infrared scanner (LI-COR). Statistical tests were performed using Prism version 7 (GraphPad).

For comparison of OROV glycoproteins from infection versus transfection, HEK293T cells were seeded at 5×10^6^ cells per 9 cm dish and allowed to rest for 24 h. Cells were transfected by mixing 20 µg of DNA (split evenly by mass between the plasmids indicated) and lipofectamine 2000 reagent according manufacture’s protocol. Alternatively, cells were infected with OROV (MOI = 1) in DMEM with 2% FCS for 1 h at 4°C on a rocker. Media was then refreshed with DMEM containing 2% FCS (infection) or 10% FCS (transfection). 24 h after transfection or infection, cell monolayers were rinsed with ice-cold PBS and then lysed on the dish with 1 mL of lysis buffer (0.1% [v/v] IGEPAL CA-630 in PBS with protease inhibitor cocktail [Sigma-Aldrich]) for 20 minutes on ice. Lysates were clarified by centrifugation 16000×g for 15 min at 4°C. The protein concentration in each lysate sample was quantified with Quick Start Bradford 1× Dye (BioRad) to equalise protein concentrations across the samples before SDS-PAGE and immunoblotting.

For glycoprotein isolation following drug treatment, HEK293T cells were transfected with a plasmid expressing HA-tagged M polyprotein as described above. After 18 h, the culture medium was collected and glycoproteins were isolated by ultracentrifugation as described above (“Before drugs”). The medium was replaced with FreeStyle 293 medium (ThermoFisher) supplemented with 5 µg/mL brefeldin A (Enzo Life Science BML-G405) or 1 µM monensin (Merck M5273) and cells were incubated for a further 6 h before the supernatant and cell lysates were harvested and processed as described above (“After drugs”).

For glycoprotein isolation from ecdysone-responsive stable cell lines, cells were seeded differently. Parental and GFP-VPS4wt EcR-293 cells were seeded at 5×10^6^ cells per 9 cm dish, whereas GFP-VPS4E/Q EcR293 cells (which grow more slowly) were seeded at 8×10^6^ cells per 9 cm dish. Cells were transfected with 7.7 µg of a plasmid expressing HA-tagged M polyprotein using TransIT-LT1 (Mirus) and after 6 h the cells were treated with either DMSO or 1 µM ponasterone A (Pon A). After 18 h of drug treatment, the medium was refreshed using FreeStyle 293 medium (ThermoFisher). Cell lysates and supernatants were harvested at 48 h post-transfection and processed as described above.

For OROV infection of ecdysone-responsive stable cells lines, EcR-293 cells were seeded as described above and infected with OROV (MOI = 1) in DMEM with 2% FCS for 1 h at 4°C on a rocker. Media was then refreshed (DMEM with 2% FCS). After 4 h cells were treated with 1 µM Pon A (or DMSO as a control) and after 24 h cells and supernatant were harvested as described previously (6).

### Co-immunoprecipitation of transfected cells

HEK 293T cells were seeded at 5×10^6^ cells per 9 cm dish. Cells were transfected with 7.7 µg of a plasmid (split evenly by mass between the plasmids indicated) using TransIT-LT1 (Mirus) and after 18 h the cells were mock-treated (refreshed media) or treated with 1 µM monensin for 6 h. Cells were pelleted (220 g, 5 min, 4°C), washed three times with cold PBS, and cell pellets were processed for immunoprecipitation.

For GFP capture, cells were lysed at 4°C in 1 mL lysis buffer (10 mM Tris pH 7.5, 150 mM NaCl, 0.5 mM EDTA, 0.5% IGEPAL CA-630, EDTA-free protease inhibitor cocktail [Roche]) for 30 min before clarification (20,000×g, 10 min, 4°C). For HA capture, cells were lysed at 4°C in 1 mL lysis buffer (10 mM Tris pH 7.5, 150 mM NaCl, 0.1% IGEPAL CA-630, EDTA-free protease inhibitor cocktail [Roche]). Protein concentration in the lysates was quantified by BCA assay (ThermoFisher) to equalise protein concentrations across the samples before immunoprecipitation with GFP-Trap resin (gta-20, ChromoTek) or anti-HA affinity matrix (11815016001, Roche) following the manufacturers’ protocols. Cell lysate was immunoprecipitated for 2 h with GFP-TRAP or overnight with anti-HA affinity matrix, at 4°C under agitation. Input samples were collected prior to immunoprecipitation and represent 2% of the total cell extract. Samples were eluted by incubation at 95°C for 5 min in 45 μL 2× SDS-PAGE loading buffer. Input and bound samples were separated by SDS-PAGE and analysed by immunoblot.

### Immunocytochemistry

HeLa cells were seeded on glass coverslips at a density of 5×10^4^ cells per well in 24 well dishes. Cells were transfected with 250 ng of DNA (split evenly by mass between the plasmids indicated) using TransIT-LT1 (Mirus). Cells were either incubated for 24 h (untreated), or for 18 h whereupon the culture medium was replenished with medium 1 µM monensin, 5 ug/mL brefeldin A or DMSO as a control before incubation for a further 6 h. Cells were transferred onto ice and were washed with ice-cold PBS and fixed with cold 250 mM HEPES pH 7.5, 4% (v/v) electron microscopy-grade formaldehyde (PFA, Polysciences) for 10 min on ice before and then incubated with 20 mM HEPES pH 7.5, 4% (v/v) PFA at room temperature for a further 20 min. After washing with PBS, cells were permeabilized by incubation with 0.1% saponin in PBS for 30 min before being incubated with blocking buffer (5% [v/v] FBS, 0.1% saponin in PBS) for 30 min. Primary antibodies were diluted in blocking buffer and incubated with coverslips for 2 h. Coverslips were washed five times with blocking buffer before incubation for 1 h with the relevant secondary antibodies diluted in blocking buffer. Coverslips were washed five times with blocking buffer, three times with 0.1% saponin in PBS, three times with PBS, and finally with ultrapure water. Coverslips were mounted using Mowiol 4-88 (Merck) containing 200 nM 4′,6-diamidino-2-phenylindole (DAPI) and allowed to set overnight. Cells were analysed on a Zeiss confocal laser scanning microscope (LSM)780 (Zeiss) or on an Olympus IX81 wide-field fluorescence microscope. Images were processed using Fiji (48, 49).

Ecdysone-responsive stable cell lines were seeded on poly-d-lysine treated glass coverslips at a density of 2×10^4^ cells per well in 24 well dishes. Cells were transfected with 250 ng of plasmid expressing HA-tagged M polyprotein using TransIT-LT1 (Mirus) and after 6 h cells were treated with 1 µM Pon A or DMSO as a control. After 18 h of drug treatment, cells were processed as described above.

## Acknowledgements

The authors thank Susanna Colaco for superb technical assistance. This work was supported by doctoral studentship and travelling fellowship from the Fundação de Amparo à Pesquisa do Estado de São Paulo (FAPESP; 2016/18356-4 and 2019/02945-9) to NSB, by a FAPESP Pump Priming award (2019/02418-9) to LLPdS, by a Biotechnology and Biological Sciences Research Council (BBSRC)/FAPESP pump priming award (BB/S018670/1) to CMC, LLPdS and SCG, and by a Sir Henry Dale Fellowship co-funded by the Royal Society and the Wellcome Trust (098406/Z/12/B) to SCG. The funders had no role in study design, data collection and analysis, decision to publish, or preparation of the manuscript. For the purpose of open access, the authors have applied a Creative Commons Attribution (CC BY) licence to any Author Accepted Manuscript version arising from this submission.

## Conflict of Interest

The authors declare they have no conflict of interest.

## Notes

### Competing Interest Statement

The authors have declared no competing interest.

### Summary of Updates

Extensively updated and improved in response to reviewer comments

